# Immune response modulation by *Pseudomonas aeruginosa* persister cells

**DOI:** 10.1101/2023.01.07.523056

**Authors:** Cody James Hastings, Grace Elizabeth Himmler, Arpeet Patel, Cláudia Nogueira Hora Marques

## Abstract

Bacterial persister cells – a metabolically dormant subpopulation tolerant to antimicrobials – contribute to chronic infections and are thought to evade host immunity. In this work, we studied the ability of *Pseudomonas aeruginosa* persister cells to withstand host innate immunity. We found that persister cells resist MAC-mediated killing by the complement system despite being bound by complement protein C3b at levels similar to regular vegetative cells, in part due to reduced bound C5b - and are engulfed at a lower rate (10-100 fold), even following opsonization. Once engulfed, persister cells resist killing and, contrary to regular vegetative cells which induce a M1 favored (CD80+/CD86+/CD206-, high levels of CXCL-8, IL-6, and TNF-α) macrophage polarization, they initially induce a M2 favored macrophage polarization (CD80+/CD86+/CD206+, high levels of IL-10, and intermediate levels of CXCL-8, IL-6, and TNF-α), which is skewed towards M1 favored polarization (high levels of CXCL-8 and IL-6, lower levels of IL-10) by 24 hours of infection, once persister cells awaken. Overall, our findings further establish the ability of persister cells to evade the innate host response and to contribute chronic infections.

## Introduction

Recurrent and persistent bacterial infections, chronic infections, are of major importance across the world as they remain in the host for extended periods of time and as such, are detrimental to patients and a burden on the healthcare system. The recalcitrance of chronic infections is aided by the characteristics of persister cells. Persister cells are present in most bacterial species, and depending on the bacterial growth phase, they can make up to 1% percent of the overall bacterial population (1, 2). Persister cells are characterized by their tolerance to various stresses, including antimicrobial agents and their formation has been attributed to antibiotic use (3), inoculum conditions and stage of growth (4), toxin-antitoxin (TA) module expression (5, 6), induction of dormancy following the stringent response and increased abundance of polyP compounds (7), deactivation of metabolism and protein synthesis genes upon exposure to fluroquinolones (8, 9), and inhibition of protein synthesis (10). Once the stress situation is removed and a carbon source or a signaling molecule indicating appropriate growth conditions are present, persister cells can revert into an antibiotic susceptible, active state (11, 12). The characteristics of persister cells render the use of most antimicrobials for the treatment of chronic infections ineffective (13).

Despite the possible contribution to chronic bacterial infections, the interactions between bacterial persister cells and the host innate immune system are still poorly understood. Typically, during an infection, macrophages are the first immune cells to recognize pathogens via their pattern recognition receptors (14). Macrophages can typically polarize toward a M1 type or M2 type response to meet the needs of the host. In an active infection, a M1 response is typically detected and is characterized by the presence of the macrophage cell membrane marker CD80, the production of reactive oxygen species (ROS) and pro-inflammatory cytokines such as IFN-β, IL-1, IL-6, IL-12, and TNF-α (15). M1-polarized macrophages typically promote differentiation of Th1 and Th17 T cells, which create an inflammatory feedback loop ultimately leading to the clearance of pathogens (16). However, it has previously been demonstrated that during infections with biofilms a M2 response is mostly detected where the cells contain the cell membrane marker CD206 and secrete anti-inflammatory cytokines, such as IL-10 and CCL5 (15). M2-polarized macrophages include a subdivision known as regulatory macrophages, or M2b macrophages - these cells play a major role in modulating the inflammatory response and are distinguished by secretion of TNF-α and IL-6, high IL-10 secretion combined with low IL-12, and expression of cell marker CD86 (17).

The complement system functions as a nonspecific defense against invading bacteria, particularly a defense against Gram-negative bacteria. Several mechanisms of complement resistance by various species of Gram-negative bacteria have been previously reported including: capsular modulation which can conceal antibody epitopes (18), the decrease of complement-mediated phagocytosis (19), resistance to the insertion of the terminal complex of complement proteins C5b-C9 in the bacterial membrane resulting in an absence of the membrane attack complex (MAC) (20, 21), and by sialyation of lipooligosaccharides on their surface (22, 23). These findings have yet to be confirmed for persister cell populations. Although little is known regarding the innate immune response to persister cells, several studies found that persister cells can survive inside macrophages (24–26) and are engulfed at a lower rate following infections (27, 28). It has also previously been found that the immune system can induce a persister state in several bacterial species such as when *S. aureus* is exposed to host oxidative stress (29), when *Salmonella* spp. are internalized by macrophages (24), upon exposure of *Mycobacterium tuberculosis* to cytokines (25), and when *Vibrio splendidus* is exposed to host-derived sea cucumber coelomic fluid (30).

*Pseudomonas aeruginosa* contributes to lung infection in cystic fibrosis patients (CF) (31) and to chronic obstructive pulmonary disease (COPD) (32), and has simultaneously been described as a one of the model systems for researching persister cells (33, 34). Furthermore, clinical isolates of *P. aeruginosa* from CF patients have been identified as high-persistence strains (35). As such, we sought to determine some of the mechanisms which enable persister cells to evade or modulate the innate immune response.

## Results

### The cell size of the P. aeruginosa persister cell subpopulation is more homogeneous than that of the P. aeruginosa vegetative population

Persister cells are a sub-population of the regular vegetative bacterial population. As such, we initiated this work by determining whether there is a substantial difference in cell size between the vegetative population of *P. aeruginosa* and isolated persister cells. Persister cells were isolated using ciprofloxacin exposure for a period of 24 h where a biphasic killing was present and cells had an exponential killing in the first 3 h followed by a plateau from that point onwards, as previously described (3, 12, 27, 33, 36, 37). To ensure that no dead cells were present, we performed several centrifugation/washing steps and the resulting pellet at the end was significantly reduced in total cell counts, as expected as, the dead cells were removed (Fig. S1). In addition, we have also performed live/dead staining using SYTO9 (stains all cells) and propidium iodide (stains cells with impaired membranes) to determine whether the final pellet of persister cells and regular cells contained similar ratios of live/dead (Fig 1A and Fig. S1B,C). We found that no significant difference was detected between the persister, and regular cell populations and that the dead cells’ control (ethanol treated cells) had more than 90% of dead cells. While overall there was no significant difference (P>0.05) in the median forward-scattering (cell size) or side-scattering (cell granulation) between regular vegetative cells and persisters, the latter had a narrower peak, indicating a less heterogeneous population, regarding cell size (Fig. 1B - F), consistent with the fact that cells, while in the persister state, are not dividing. However, these cells have been shown to be phenotypically different in *E. coli* (38–40).

**Figure 1.**
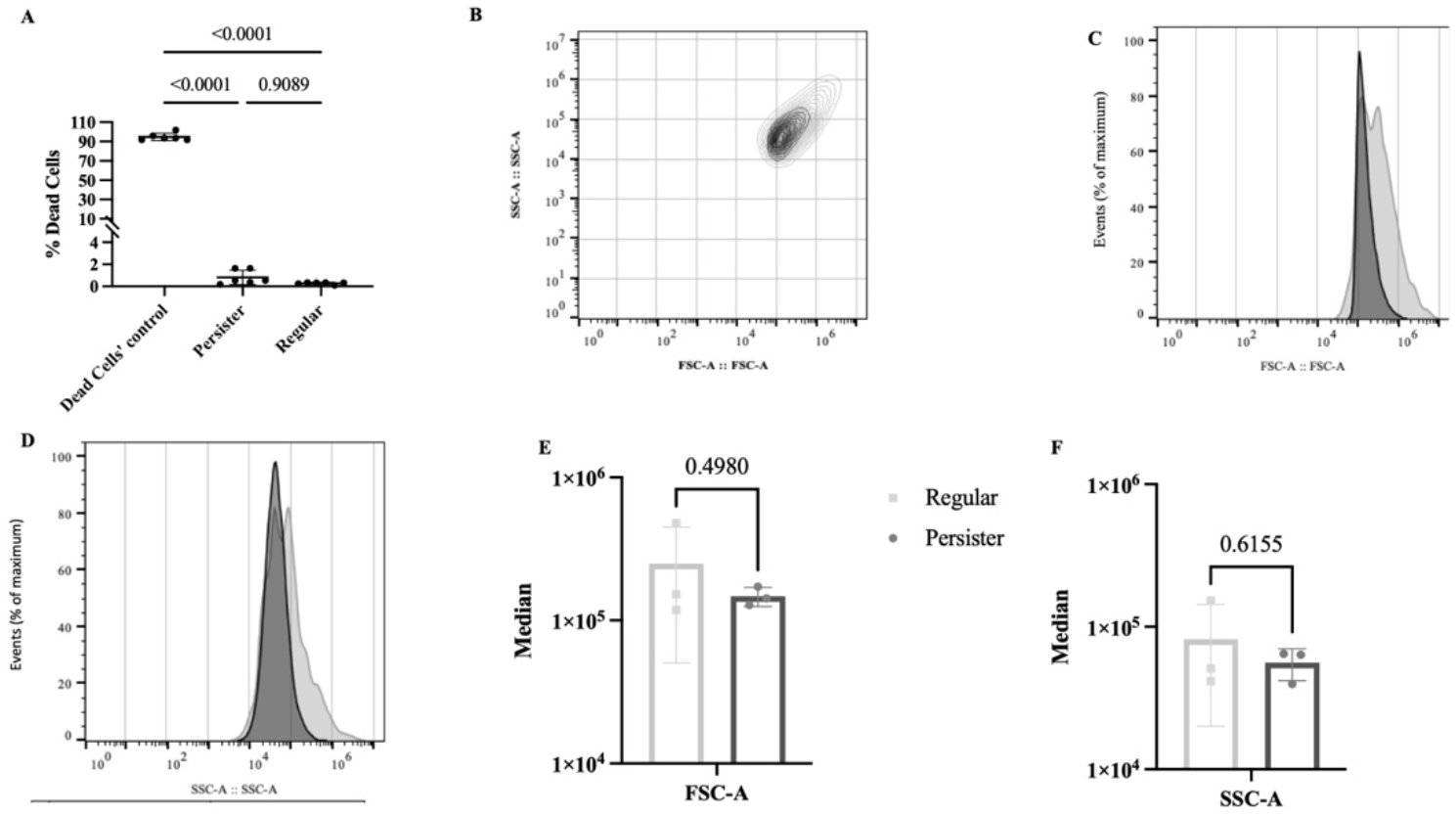
Live/dead cell ratio and cell sizes. Regular and persister cells of *P. aeruginosa* were isolated and then stained with Syto9/propidium iodide to determine live/dead ratio and with BacLight red for FACS analysis. A. Live/Dead, B. Size cell scattering, C. Forward scattering, D. Side scattering, E. Median of the Forward scattering peak size, F. Median of the Side scattering peak size. Results were analyzed with T-test (*P<0.05) and are presented as mean ± SD.

**Figure S1.**
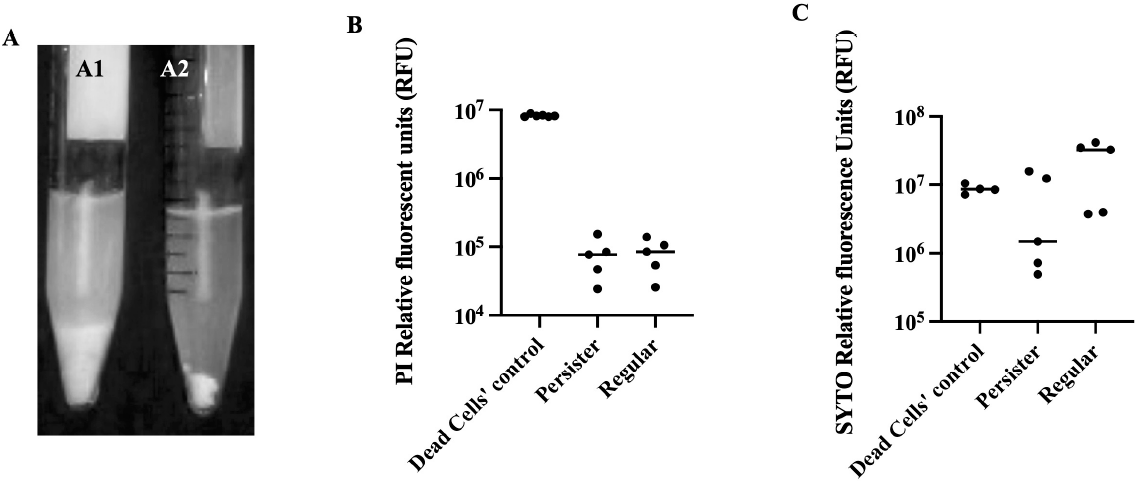
Cells remaining following selection. Regular vegetative cells and persister cells of *P. aeruginosa* PA14 were isolated from stationary phase planktonic cultures. Cells exposed to saline were named regular vegetative cells (Al) and cells exposed to ciprofloxacin (20 mg/L) in saline were named persister cells (A2). Subsequently, isolated/selected cells were stained with SYTO9 and propidium iodide and the % of dead cells was calculated (B).

### P. aeruginosa persister cells are tolerant to MAC-mediated killing despite being opsonized by C3b

We next determined the interaction of *P. aeruginosa* persister cells with human complement both for MAC-mediated killing (Fig. 2) and opsonization (Fig. 3). Due to its ability to inhibit and resist MAC-mediated killing (41, 42), *Staphylococcus aureus* was used as a negative control (Fig. 2A-B), whilst due to its susceptibility to MAC-mediated killing (43), *Escherichia coli* was used as a positive control (Fig. 2C-D). In the presence of serum with inactivated complement, persister cells of each species reverted to an active dividing state (Fig. 2 B, D, F). As anticipated, *S. aureus* viability of both regular vegetative (Fig. 2A) and persister (Fig. 2B) cells was unaffected by the presence of complement (Fig. 2A-B) while viability of *E. coli* regular vegetative cells was reduced to the point of eradication (Fig. 2C). Contrary to what was previously described (43), *P. aeruginosa* regular vegetative cells (Fig. 2E) were eradicated, albeit at a lower rate initially when compared to *E. coli* regular vegetative cells (Fig. 2C). Both *E. coli* (Fig. 2D) and *P. aeruginosa* (Fig. 2F) persister cells were initially killed at an increased rate but by 1.5-3 h presented a biphasic killing trend, becoming resilient to killing by complement.

**Figure 2.**
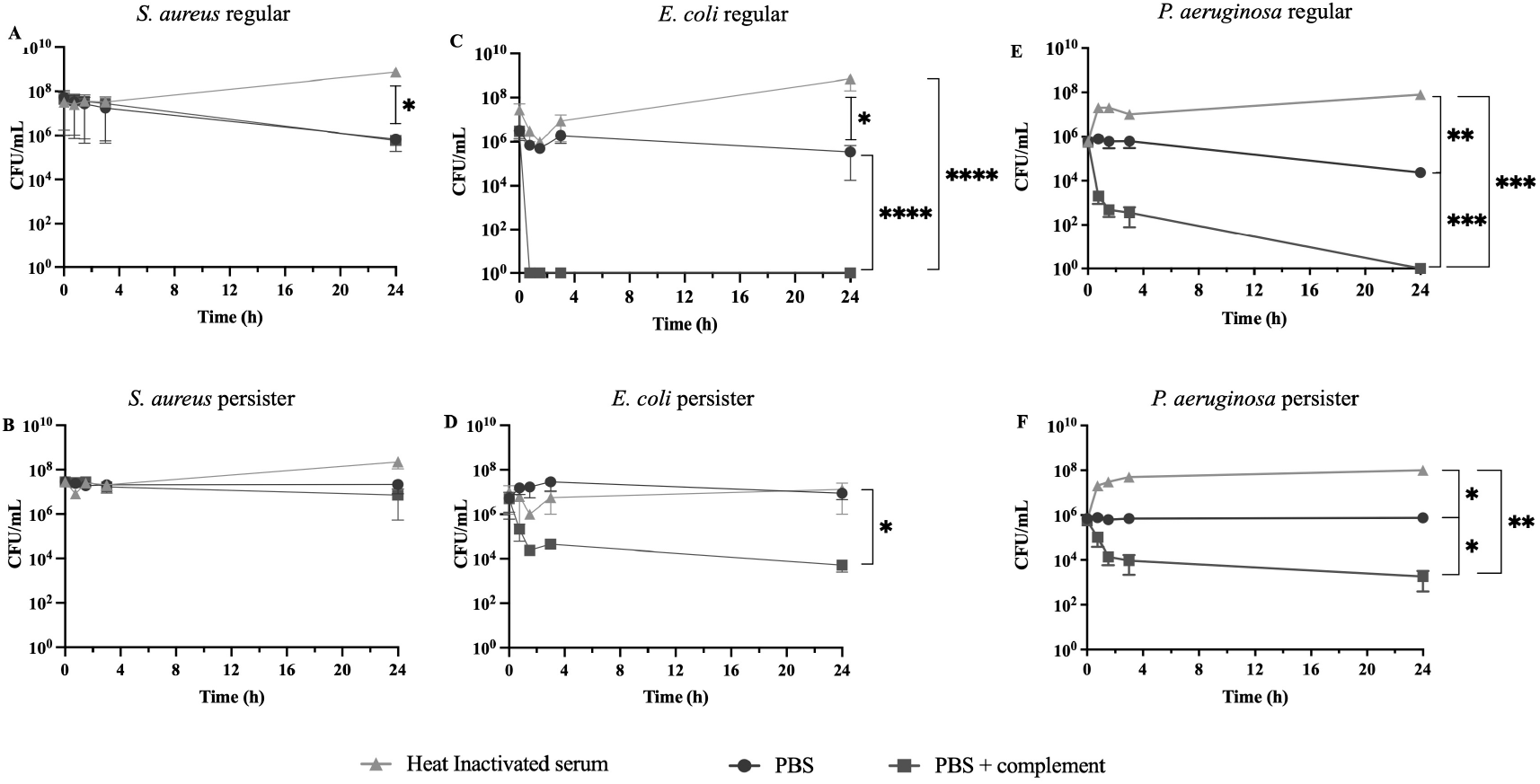
Time kill curves of complement factor proteins. Regular and persister cells of *S. aureus* (A, B), *E. coli* (C, D), and *P aeruginosa* (E, F) were exposed to 90% complete human serum (PBS+complement) (closed square), PBS(closed circle), or heat inactivated serum (closed triangle) for a period of 24 h. Experiments were performed in quadruplicate. Results were analyzed according to the one-way ANOVA with Tukey’s post-test (*P<0.05, **P<0.01, ***P<0.001) and are presented as mean ± SD.

**Figure 3.**
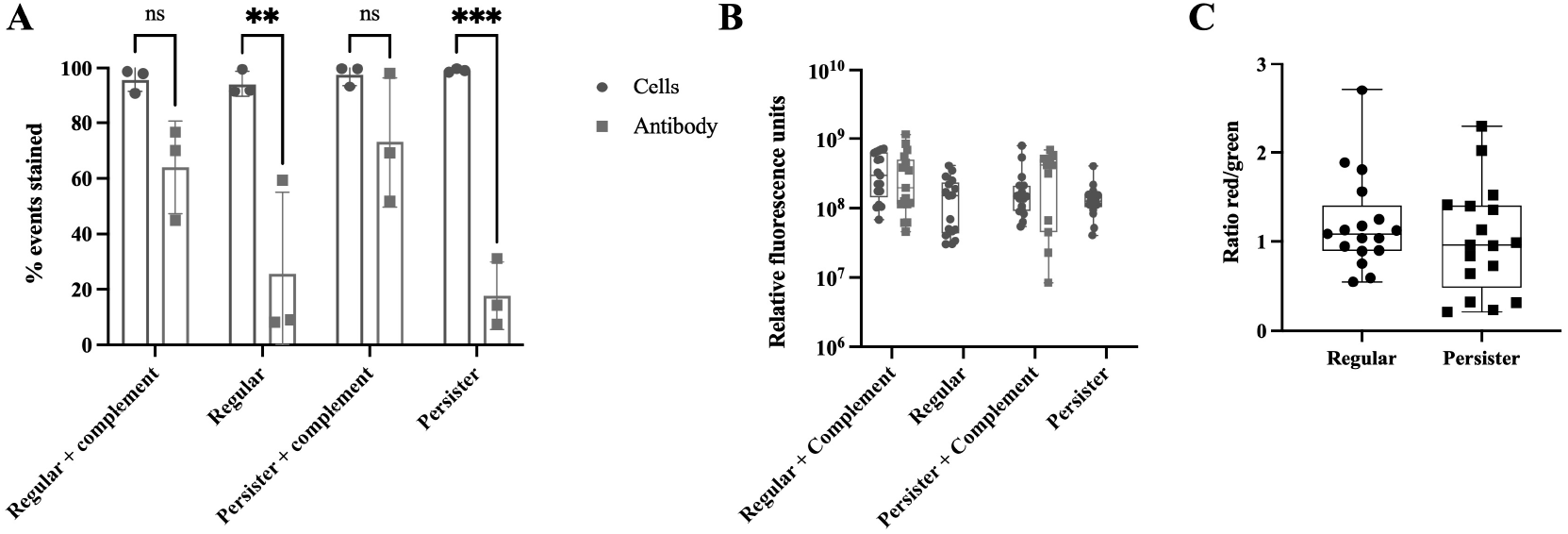
Binding of complement factor C3b to cells. Persister and regular cells were isolated and exposed to human serum containing complement for a period of 30 min. Upon ending the complement reaction with EDTA, the cells were stained with Baclight red and complement was immunostained (green) and cells were sorted via FACS (**A**) or images of the cells were acquired via epifluorescence microscopy: **B** represents the relative fluorescence of red stain and green stain in regular and persister cells upon exposure to complement, **C** represents the ratio of cells to antibody, when analyzing fluorescence images using the Luminance program. A total of 3 experiments were performed where 20 images were used in this experiment and analyzed using the Intensity Luminance Software. Results presented as mean ± SD. When performing image analysis, no statistical difference was found between the staining of regular vs persister cells, as determined by one-way Anova. When performing FACS, a significant difference was found between cells and antibody, in the absence of complement as determined by two-way ANOVA with Tukey’s post-test (**P<0.01, ***P<0.001)

### P. aeruginosa persister cells are opsonized by C3b similarly to regular vegetative cells but have reduced bound C5b

To establish whether the resilience of *P. aeruginosa* to complement killing was due to an inability of C3b (initiating protein of the alternate complement pathway) and/or C5b (initiating protein of the MAC formation) - to bind persister cells we used an anti-C3 antibody to detect and quantify the C3b binding to the cells (Fig. 3) and performed ELISA for C5b protein quantification (Fig. 4). The binding quantification was performed by FACS (Fig. 3A) and further confirmed by microscopy (Fig. 3B). We found no significant difference (P>0.05) of C3b binding between persister and regular vegetative cell populations albeit persister cells having a clear bi-modal pattern of binding (Fig. 3B). We also found that after 1.5 h and 3 h of incubation in human serum, significantly less C5b was deposited on the viable persister cells’ membranes relative to viable regular cells, while after 24 hours no regular cells were viable (Fig. 2E), resulting in much more C5b deposition per viable cell in the persister population (Fig 4). No significant change in C5b deposition on persister cells was detected from 3 h onwards (P>0.05) (Fig. 4).

**Figure 4.**
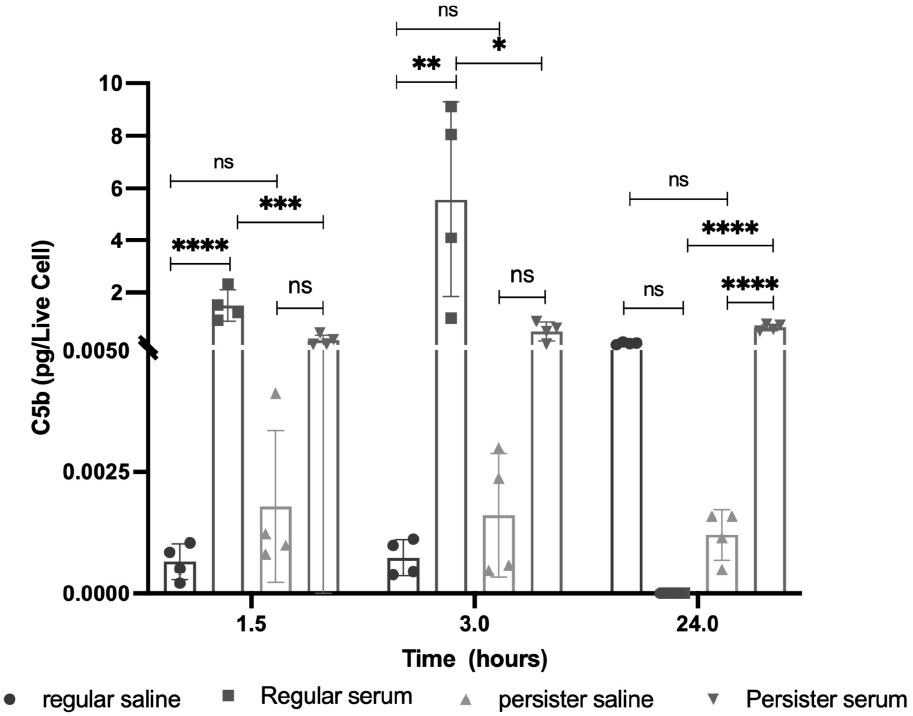
Binding of complement factor C5b to cells. Persister and regular cells were isolated and exposed to human serum containing complement for a period of 1.5, 3, and 24 hours. Upon ending the complement reaction with EDTA, the cells were harvested and the presence of C5b on the cell envelope was quantified using ELISA. Results presented as mean ± SEM. Statistical significance was determined by ANOVA with Tukey’s post-test, * P<0.05, **P<0.005, *** P<0.001, ****P<0.0001.

### Macrophages can engulf P. aeruginosa persister cells, albeit at a lower rate, but do not kill them

Upon determining that persister cells were not killed by the membrane attack complex function of complement (Fig. 2), were opsonized with C3b similarly to regular vegetative cells (Fig. 3), and had lower C5b bound (Fig. 4), we quantified the macrophages’ ability to engulf *P. aeruginosa* persister cells with and without prior-opsonization. THP-1 macrophages were exposed to the same inoculum of bacteria, whether regular or persister, for 30, 60, 90, and 180 min and engulfment was evaluated based on intracellular bacterial viability (Fig. 5). Persister cells of *P. aeruginosa* were engulfed significantly less (P<0.001) by THP-1 macrophages compared to regular vegetative cells, with an overall 100-fold decrease (Fig. 5). As anticipated, opsonization of regular vegetative cells did not result in a significant change in engulfment (44). However, we expected a change to occur for persister cells but, although the engulfment was slightly higher following opsonization of persister cells, it was neither significant (P>0.05) nor to the level of regular vegetative cells (Fig. 5).

**Figure 5.**
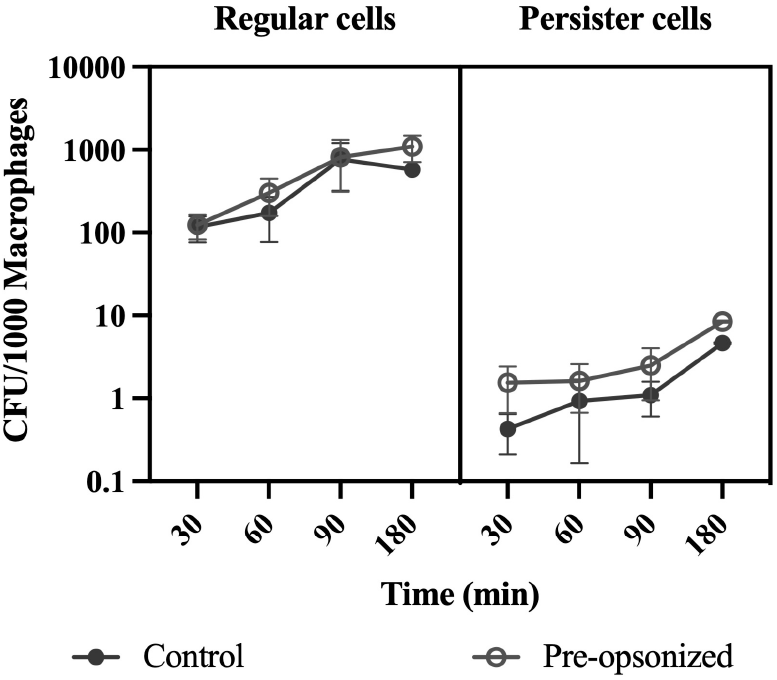
Macrophage Infections. Differentiated THP-1 monocytes were exposed to *P aeruginosa* regular and persister cells for 30, 60, 90, and 180 min. Regular cells were diluted to match viable persister cell concentrations before inoculation. Bacterial viability was quantified at the various time points of the infection. Results were analyzed according to the one-way ANOVA with Tukey’s post-test and are presented as mean ± SEM. Experiments were performed in quadruplicate. Engulfinent of persister cells was significantly lower comparing to regular cells (P<0.001) but no significant difference (P<0.05) was observed when cells were exposed to complement.

### P. aeruginosa persister cells are resilient to killing by macrophages

Typically following engulfment, the phagosome fuses with a lysosome and the engulfed pathogenic organisms are eliminated. Thus, once it was established that persister cells were resilient to killing by complement (Fig. 2) and were engulfed at a lower rate (Fig. 5) when compared to regular vegetative cells, we decided to further explore their resilience to killing once inside the macrophages. Clearance of *P. aeruginosa* persister cells was quantified by infecting macrophages for 1.5 h, subsequently removing all external bacteria, and then allowing the macrophages to kill the intracellular bacteria for a period of 24 h. The number of viable intracellular regular vegetative cells present was significantly reduced (P<0.05) by a total of 1.4 Logs (Fig. 6A), while no change of cell viability was detected for the intracellular persister cells (Fig. 6A) indicating a lack of killing by macrophages. The viable cell count at 24 h post infection was similar for infections with both regular vegetative and persister cells (Fig. 6A). FACS analysis of internalized cells within macrophages, at 90 min of infection and 24 h post infection, where bacteria were stained with Baclight red®, also showed fewer regular vegetative cells (with a shift of the fluorescence peak) but not fewer persister cells (Fig. 6B). In addition, we also quantified the 16s rRNA gene expression, to determine whether the *P. aeruginosa* cells were active in the macrophages upon engulfment and found that it was decreased at time 0 by 5.6-fold ± 1.7 and no change at time 24 (1.1 ± 0.4) in persisters compared to regular vegetative cells. These findings provide evidence that the only surviving cells within the macrophages 24 h post infection are persister cells and are supported by previous findings for *Salmonella typhimurium* where once engulfed the bacteria adopt a non-growing antibiotic tolerant state and can reside for extended periods of time within the macrophages (45).

**Figure 6.**
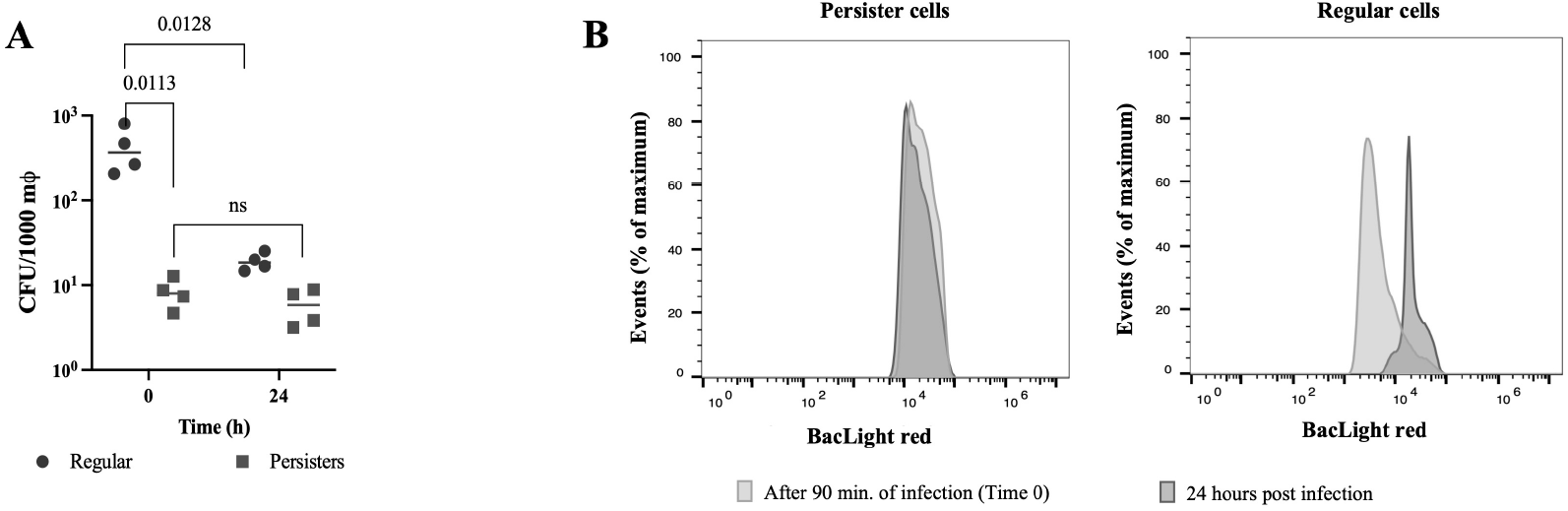
Elimination of *P. aeruginosa* cell populations 24 h post infection. THP-1 macrophages were infected with regular and persister cells (MOI of 10). After 90 min of infection, the intracellular bacteria were quantified (Time 0). Extracellular bacteria were removed with gentamicin and media was replaced. The intracellular bacteria were quantified at 24 h of incubation to determine bacterial elimination via viable counts **(A)** and FACS **(B).** Viable counts were analyzed according to the one-way ANOVA with Tukey’s post-test.

### P. aeruginosa persister cells elicit an intermediate inflammatory response and a macrophage favored a M2 polarization

It was thus clear that *P. aeruginosa* persister cells are resilient to killing by complement (Fig. 2) and by macrophages (Fig. 6), while also being engulfed at a lower rate (Fig. 5). Therefore, we questioned whether upon infection with persister cells, macrophages - main source of cytokine secretion following infection - were responding similarly to an infection by regular vegetative cells. During an infection, macrophages can typically polarize toward a M1 type or M2 type response to meet the needs of the host. (15) To establish the response to infections with persister and regular vegetative cells, we quantified the secretion (Fig. 8) and relative gene expression (Fig. S3) of the pro-inflammatory cytokines CXCL-8, IL-6 and TNF-α - indicative of a M1 response - together with the anti-inflammatory cytokine IL-10 - indicative of a M2 response (15), at 0.5, 1.5, and 24 h of infection. We also quantified the presence of CD80, CD86, and CD206 on the macrophage cells (46), using flow cytometry (Fig. 7). When quantifying the cell membrane markers, we found that after 1.5 h of infection, macrophages infected with vegetative *P. aeruginosa* cells expressed high levels of CD80/CD86, but not CD206 (Fig. 7). In contrast, macrophages infected with persister cells expressed both high levels of CD80/CD86 and CD206 (Fig. 7). These results suggest that infections with persister cells elicit macrophage polarization towards a M2 response while still retaining M1-associated cell surface proteins. Regarding cytokine secretion, we found that in the first 1.5 h, all inflammatory cytokines were secreted at lower levels by macrophages infected with persisters compared to infections by regular vegetative cells, but higher than unchallenged macrophages (Fig. 8 A-C). Similarly, the anti-inflammatory IL-10 also presented that pattern at 0.5 h; however, at 1.5 h minutes of infection, IL-10 secretion was significantly higher in persister-infected macrophages than in infections with regular vegetative cells (Fig. 8D). This high anti-inflammatory response coincides with the plateau of engulfment established between 0.5 and 1.5 h, for persister cells (Fig. 4), and the consistently lower engulfment of persister cells. However, at 24 h of infection with persister cells, a bi-modal trend was present in IL-6 (Fig. 8A) and CXCL-8 (Fig. 7C) with an overall increase in secretion, compared to 1.5 h, whilst IL-10 was at levels identical to uninfected macrophages (Fig. 8D). These changes were anticipated as, similarly to what occurs in infections *in vivo* (27), a percentage of the persister population reverted into an active metabolic state due to the presence of nutrients in the medium following 7 h of incubation, albeit at significantly lower levels than regular vegetative cells (Fig. S2), and as such, should activate the pro-inflammatory response while tampering the anti-inflammatory response as demonstrated by the decrease of IL-10. In infections with regular vegetative cells, the IL-6 (Fig. 8A) and TNF-α (Fig. 8B) inflammatory response, continued to increase in the first 1.5 h, remaining constant at 24 h, whilst CXCL-8 decreased by 24 h (Fig. 8C). IL-10 remained constant in the first 1.5 h, decreasing to uninfected control levels by 24 h (Fig. 8D).

**Figure 7.**
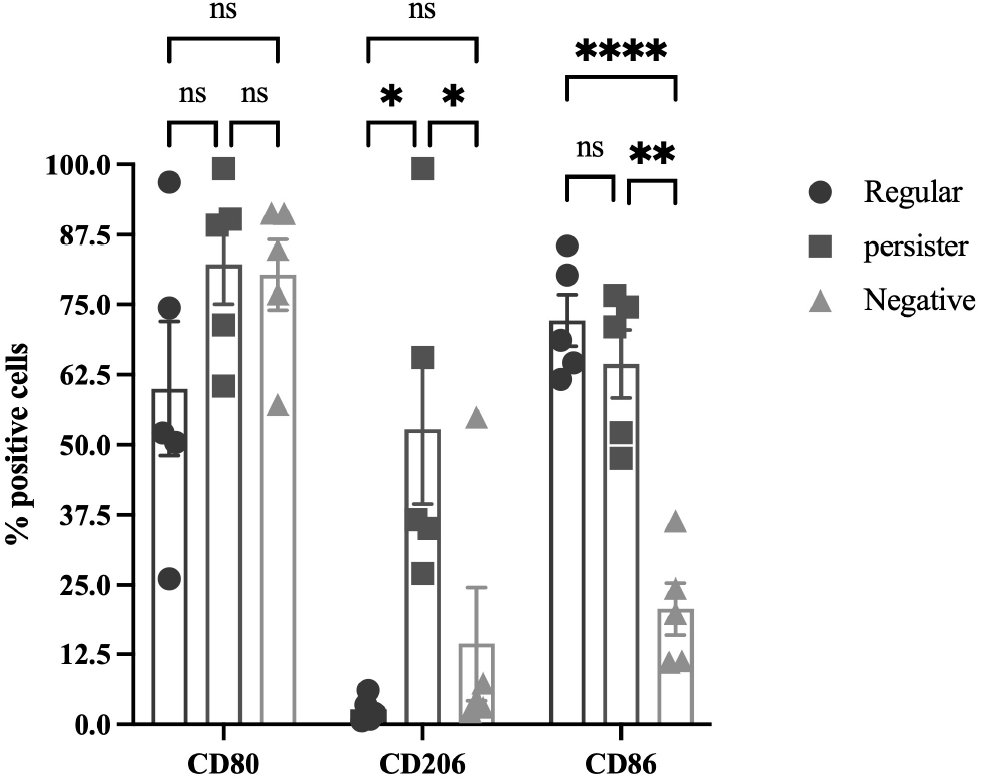
Macrophage polarization. THP-1 macrophages were infected with regular and persister cells (MOI of 10). After 90 min of infection, extracellular bacteria were removed with gentamicin then cells were trypsinized and subsequently incubated with Anti-CD8O, Anti-CD86, and Anti-CD2O6 antibodies, then washed and resuspended in PB A for FACS analysis. Results presented as mean ± SD. *P<0.05, ** P<0.005, **** P<0.0005 as determined by one-way Anova followed by a Tukey’s multicomparison test.

**Figure 8.**
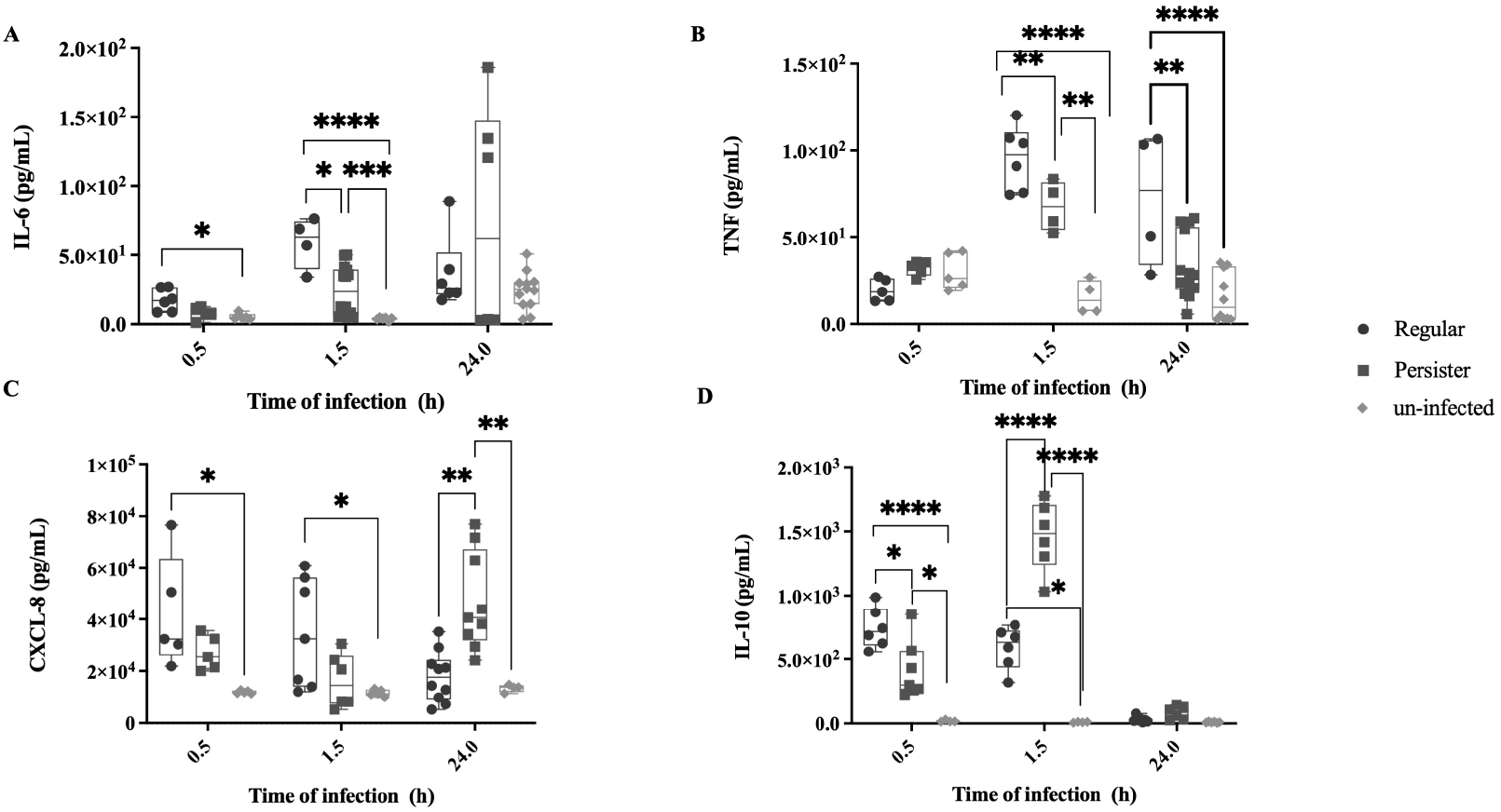
Cytokine secretion by macrophages. Cytokine secretion of macrophages was quantified for 0.5, 1.5, and 24 h of infection (MOI of 10) with *P aeruginosa* regular and persister cells, and uninfected (control). (A) IL-6, (B) TNF, (C) CXCL-8, (D) IL-10. Results shown consist of at least 4 experiments. *P<0.05, ** P<0.005, *** P<0.001, **** P<0.0005 as determined by one-way Anova followed by a Tukey’s multicomparison test.

**Figure S2.**
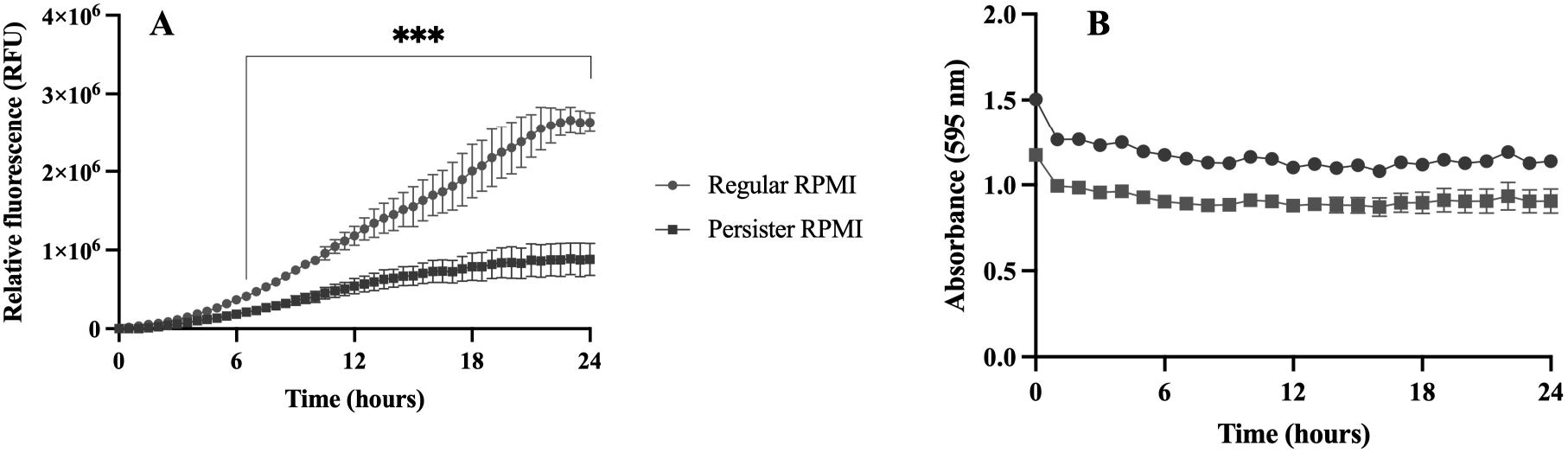
Bacterial growth and metabolism of regular vegetative and persister cell. Bacterial populations of *P. aeruginosa* PA14 and MPAOl attTn7::P(Al/04/03)::GFPmut were isolated and resuspended in RPMI medium. Constitutive fluorescence of MPAOl attTn7::P(Al/04/03)::GFPmut (A) and absorbance of PA14 (B) were monitored for 24 hours. Results were analyzed according T-test (***P<0.001) and are presented as mean ± SD.

To determine whether the variation of cytokine secretion of macrophages upon infections with persister and regular cell infections (Fig. 8) was transcriptionally regulated, we quantified the relative expression of the genes related to the several cytokines (Fig. S3). We found that except for the 24 h time point of IL-6, no significant difference in the gene expression quantification was present, indicating that the macrophage’s transcription of cytokine genes is equally activated when exposed to both cell types. As such, post-transcriptional regulation must be occurring as the cytokine secretion is significantly different between infections with regular and persister cells (Fig. 7). In infections with both bacterial populations cytokine mRNA is still transcribed, but due to the persister cells’ low metabolic status, there is a reduction/absence of microbial products which triggers the macrophage response into an event similar to the clearance of microbial products *in vivo,* where it is known that the mRNA coding for cytokines becomes unstable resulting in a reduction of translation (47) and an absence of bacterial elimination. Thus, this links the absence of microbial products to post-transcriptional control, which is normally used to prevent unwarranted cytokine production, explaining the intermediate cytokine secretion (Fig. 8) in infections with persister cells and the lack of elimination once engulfed (Fig. 6).

**Figure S3.**
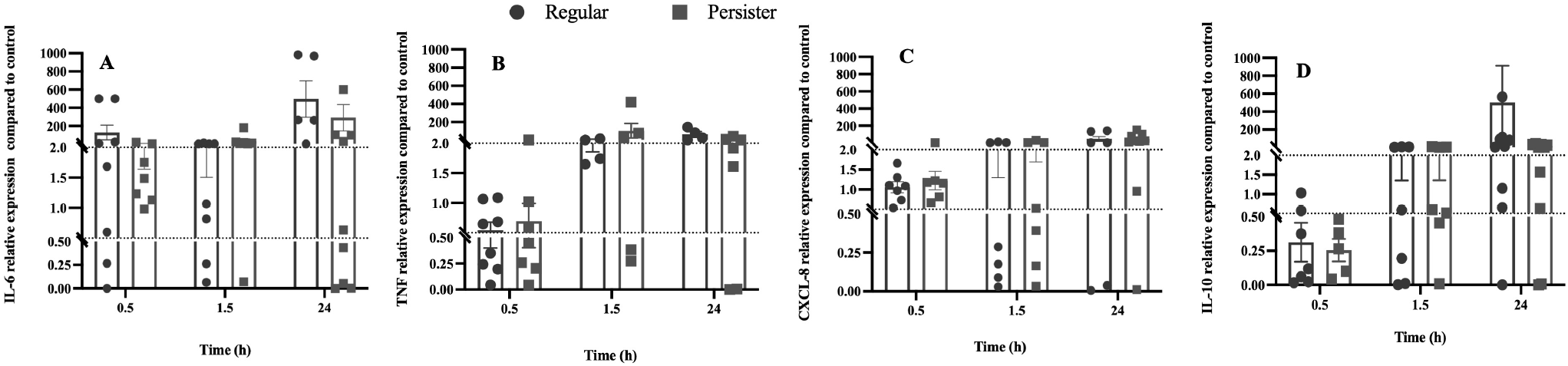
Macrophage cytokine gene expression by macrophages. The relative gene expression of 4 cytokines IL-6 (A), TNF (B), CXCL-8 (C), and IL-10 (D) was quantified for: 0.5, 1.5, and 24 h of infection (MOI of 10) with *P aeruginosa* regular and persister cells. The relative expression level for *P aeruginosa* persister and regular cells was compared to un-infected controls. Results shown consist of at least 4 experiments. The CT value of the housekeeping gene *gadph* remained constant throughout the different treatments (P>0.5 by ANOVA and no difference between treatments by Tukey’s multiple-comparison test). A significant change was considered to occur when a 2-fold change in the relative expression level occurred.

## Discussion

The innate immune response to bacterial persister cells remains ambiguous, despite their hypothesized role in chronic and resilient infections. Previously it has been found that a persister state can be induced by the immune system in several bacterial species (24, 25, 29, 30), and that persister cells are engulfed at a lower rate following infections (27, 28) and can survive once inside macrophages (24–26).

In this study, we describe a mechanism of the effect of persister cells on the immune response, whilst describing their resilience to other aspects of innate immunity namely MAC-mediated complement killing and macrophage killing. Our study provides evidence that *P. aeruginosa* persister cells (1) resist both MAC-mediated complement killing and macrophage killing albeit being opsonized by C3b, and (2) elicit the polarization of macrophages toward a M2 response, which switches to a M1 response upon persister awakening. To our knowledge, this is the first comprehensive study of the overall innate immune response to *P. aeruginosa* persister cells.

We found that there is a decreased susceptibility to MAC-mediated killing in *P. aeruginosa* and *E. coli* persister cells compared to regular - metabolically active - bacterial cells (Fig. 2). This resistance to MAC-mediated killing was due to a decrease of C5b binding (Fig. 4) but not due to a reduction of C3b deposition on the bacterial surface (Fig. 3) as previously hypothesized for the evader phenotype (43) which was most likely a subset of the persister cell phenotype as, the persister cell population in *P. aeruginosa* PA14 consists of 0.1% to 0.001% of the regular vegetative population (12), consistent with evader phenotype values (43). Functional C5 convertases have been observed on C3b-opsonized *P. aeruginosa* (48), so it is possible that these convertases are less functional on the surface of persister cells, leading to decreased deposition of C5b. Furthermore, this is in accordance with previous findings where, in *E. coli,* exposure to human serum has resulted in the induction of both the persister and the viable but non-culturable (VBNC) state (49) which has previously been shown, in *E. coli,* to describe the same bacterial stress state (50). The resilience to complement killing in both *E. coli* and *P. aeruginosa,* could be due to changes in the cell membrane and outer membrane, as previously it was reported that several outer membrane proteins (OprF, OprB, OprD, and OprM) and the chaperone protein SurA were present in higher abundance in *P. aeruginosa* cells reverting from a persister state (12). In the absence of SurA bacterial cells are highly susceptible to complement (51) whilst OprF is a complement C3 binding acceptor molecule (52), and its absence reduces the bacterial escape from phagosome vacuoles (53). A similar process could be occurring in *E. coli* as OmpA, the homolog to OprF, has been implicated in C3 convertase inhibition (52) and the inhibition of the classical complement pathway (54). It has also been established that *P. aeruginosa* can cleave complement protein C3 through binding complement Factor H via cell surface-associated proteins Tuf (Kunert et al., 2007) and LpD (55). Persister-like cells in *P. aeruginosa, E. coli,* and four other relevant human pathogens show tolerance to eradication by complement-mediated lysis, however these cells require a level of metabolic activity to persist in blood (43).

Similar to previous work that reported phagocytosis of *S. aureus* persister cells (27) and *Mycobacterium tuberculosis* (28) dormant cells was significantly lower than active/regular vegetative cells, persister cells of *P. aeruginosa* were engulfed at a lower rate when compared to regular vegetative cells. However, this was independent of bacterial cell opsonization (Fig. 3). Once engulfed, *P. aeruginosa* persister cells numbers remained constant, indicating a lack of killing, contrary to regular vegetative cells where the viable cell level was reduced to persister cell levels (Fig. 6). As such, it seems that *P. aeruginosa* switches to a persister state once inside the macrophages, similarly to the intracellular pathogen *Listeria monocytogenes* which switches to a persistence phenotype when found in vacuoles (26). However, the fate of persister cells after engulfment remains mostly unclear, and further studies need to be performed, as both *S. typhimurium* and *M. tuberculosis* persister cells are metabolically active following engulfment by macrophages (24, 25). Furthermore, *P. aeruginosa* uses Type III secretion system (T3SS) proteins to attack host phagocytes (56), and these proteins have been shown to accumulate in *P. aeruginosa* persister cells, killing host immune cells (57). We did not however, detect changes in the macrophage numbers post infection (data not shown).

When examining transcription of several cytokines characterized in M1 and M2 polarizations (Fig. S3), we found that infections with both persister and regular vegetative cells result in a similar gene transcription supporting the hypothesis that both cell populations activate the immune system, albeit with post-transcription or translation modifications. The killing of the regular vegetative cells of *P. aeruginosa* (Fig. 6) indicates that macrophages are activated upon infection, as further evidenced by them being CD80+/CD86+CD206- (Fig. 7) together with the secretion of high levels of CXCL-8, IL-6, and TNF-α, and were not deterred by the initial high concentration of IL-10 (Fig. 8D), compared to un-infected cells. In contrast, in infections with persister cells macrophages were CD80+/CD86+/CD206+ (Fig. 7) and were initially intermediately activated when exposed to persister cells, as shown by their secretion levels of IL-6, CXCL-8 and TNF-α, compared to uninfected and regular-infected macrophages, followed by a tampering down of the pro-inflammatory response, due to the high IL-10, resulting in a lack of elimination of the intracellular bacteria when the infection is stopped at 1.5 h (Fig. 6), previous to persister cell reversion to an active state in RPMI medium (Fig. S2) - as demonstrated to occur in when exposed to heat-inactivated serum (Fig. 2F), and known to occur upon the removal of stress (1, 2, 45). These results are supported by previous findings described for *S. typhimurium* persisters (45) where it was found that *S. typhimurium* persisters induced anti-inflammatory polarization of macrophages and extended the survival of the bacteria within the host (45). Additionally, when *M. tuberculosis* chronically infects the lungs of wild type mice a subpopulation of dormant cells is present, whereas mice lacking in interferon-γ lack this subpopulation (25) suggesting that the presence of host cytokines is important to the persistence of *M. tuberculosis* during infection.

From our findings, we propose that the mechanism of infection and immune system modulation between regular vegetative cells and persister cells is distinct (Fig. 9). Regular vegetative cells induce a macrophage favored polarization toward M1 (CD80+/CD86+/CD206-, high levels of CXCL-8, IL-6, and TNF-α), whilst persister cells initially induce a polarization favoring M2 - more specifically M2b (CD80+/CD86+/CD206+, high levels of IL-10, and intermediate levels of CXCL-8, IL-6, and TNF-α), which is then skewed towards M1 polarization, by 24 h of infection, once the internalized persister cells revert into an awakened metabolically active state.

**Figure 9.**
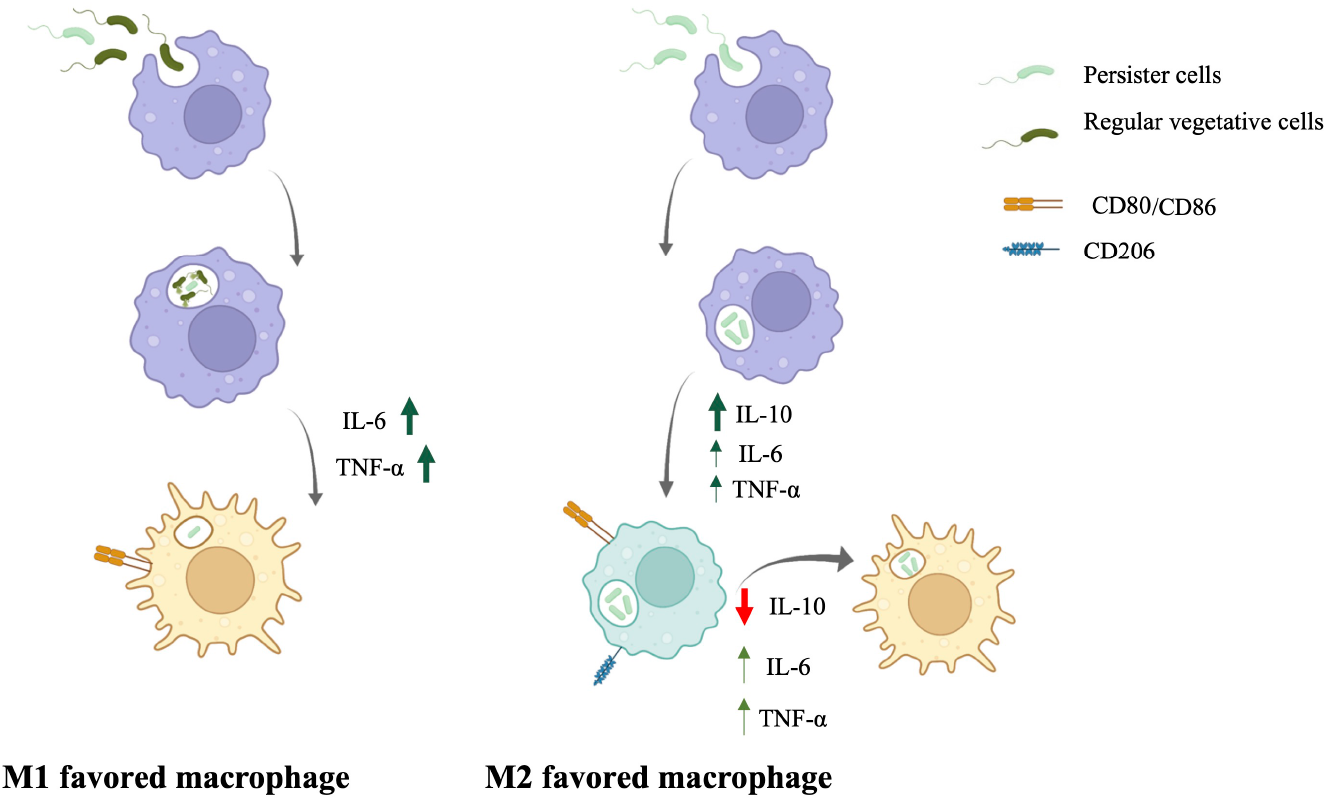
Mechanism of infection of macrophages and the modulation of the immune response. When regular vegetative cells are engulfed, macrophages polarize toward M1 and the bacterial cells are killed with only persister cells remaining. When persister cells are engulfed, macrophages polarize toward M2 (more specifically M2b) which is then skewed towards Ml polarization, once persister cells revert into an awakened metabolically active state.

In conclusion, we found that in addition to being tolerant to antibiotics, persister cells are also resilient to an immune system attack/response where they are not eliminated by MAC-mediated killing, as demonstrated by the decrease of bound C5b, despite being bound by C3b, and elicit an intermediate anti-inflammatory response by triggering macrophage M2b favored polarization. This study sheds further light as to how persister cells modulate the immune response and survive in the host during an infection. By escaping/resisting the immune response, persister cells can later become active dividing cells and re-infect the host, further confirming that these cells are involved in chronic and recurrent infections. Despite these advances, there remain many unknowns relating to persister cell behavior when infecting a host, and how the immune response to persister cells occurs in other bacterial species.

## Acknowledgements

We would like to acknowledge the support from Binghamton University structural funds, NIH educational grant 2 R25GM056637-14, and NSF grant 1139772.

## Author contributions

CNHM. Conceived the concept. CJH, GEH, and AP carried out the experiments. CNHM and CJH analyzed the data and co-wrote the paper. All authors discussed the results and commented on the manuscript.

## Methods

### Bacterial strains and growth conditions

In this study, *Staphylococcus aureus* ATCC 6538, *Pseudomonas aeruginosa* PA14, *P. aeruginosa* MPAO1 attTn7::P(A1/04/03)::GFPmut, and *Escherichia coli* BW25113 were used. Overnight cultures were grown on Lennox media (LB, Becton, Dickinson, Sparks, MD) in Erlenmeyer flasks at 37°C with aeration.

### Isolation of persister cells

Persister cells were isolated as described previously 1–6. Briefly, isolation streak plates of *S. aureus, P. aeruginosa* and *E. coli* were prepared on 100% LB agar and incubated at 37°C for 24 h. Planktonic overnight cultures were prepared by removing an isolated colony from the streak plate and inoculating it into 100% LB broth and grown at 37°C with agitation (220 rpm) for a period of 24 h. Cells were then collected (16,000 x g for 5 min at 4°C), washed twice with saline (16,000 x g for 5 min at 4°C), and subsequently resuspended in either saline (0.85% NaCl) or ciprofloxacin (20x MIC) in saline to a final OD600 of 1.6. Ciprofloxacin was used as means to induce oxidative DNA damage which results in an accumulation of persister cells (39). Cultures were subsequently incubated at 37°C with agitation (220 rpm) for a period of 24 h. Bacterial cells were collected via centrifugation (16,000 x g for 5 min at 4°C). The cells were then resuspended in saline and washed two times by centrifugation (16,000 x g for 5 min at 4°C). These 2 washes were performed to remove lysed dead cells, as ciprofloxacin was previously demonstrated to lyse cells of *P. aeruginosa, Escherichia coli,* and *Enterobacter cloacae* (58–60). Once the first wash is performed, a large amount of biomass – the dead lysed cells – is removed (Fig. S1) and only live cells were present in the final resuspension. Ciprofloxacin concentrations used were 20x the MIC and consisted of 50 mg/L for *S. aureus* cells and 20 mg/L for *E. coli* and *P. aeruginosa* (3, 12, 24, 30, 33, 34). Viability of persister and regular vegetative cells was determined at 0, 1, 3, 6, and 24 h, by serial dilution and drop plating on 1:2 plate count agar (PCA) with 1%MgCl2.7H2O for the inactivation of ciprofloxacin. Cell viability was also determined by staining persister and regular cells with propidium iodide and SYTO9 (ThermoFisher), where after a 15 min incubation, the cells were washed (to remove excess of stain) with PBS and resuspended. Bacterial fluorescence (from the stains) was measured using a SpectraMax I3x Multi-Mode plate reader, Molecular Devices. We also used dead cells – cells exposed to 70% ethanol for 30 min – as a control.

### Growth of *Pseudomonas aeruginosa* in RPMI-1640

*P. aeruginosa* regular and persister cells were isolated as above, collected by centrifugation, and washed in 0.85% saline three times by centrifugation (16000 *x g* for 5 min at 4°C). Each population was then resuspended in RPMI at a final OD_600_ of 1.5. Each population was added to a 96-well plate and changes in absorbance (595nm) was monitored every hour, for 24 hours in a microtiter plate reader (Beckman DTX880) at 37°C.

### Activation of metabolism in *Pseudomonas aeruginosa*

in RPMI-1640. Regular and persister cells of *P. aeruginosa* MPAO1 *attTn7::P(A1/04/03)::GFPmut* (61), constitutively expressing GFP, were isolated as above, collected by centrifugation, and washed in 0.85% saline three times by centrifugation (16000 *x g* for 5 min at 4°C). Each population was then resuspended in RPMI to a final OD600 of 1.5. Each population was added to a 96-well plate and changes in fluorescence (excitation 488 nm, emission 509 nm) were monitored for 24 h, at 30 min intervals (SpectraMax I3x Multi-Mode plate reader, Molecular Devices).

### Effect of Human Serum on regular and persister cells

Regular vegetative and persister cells of *P. aeruginosa, S. aureus, and E. coli* were collected by centrifugation, washed in 0.85% saline three times by centrifugation (16000 *x g* for 5 min at 4°C). Samples were then resuspended in 0.85% saline at a final concentration of 10^7^ cells/mL, and subsequently in either 90% complete human serum, PBS, or heat inactivated serum. Cells were then incubated at 37°C, and cell viability was determined by adding 10 μL of 10 mM EDTA to stop the reaction at 0, 0.75, 1.5, 3, and 24 h of incubation followed by serial dilutions and plating of bacteria on 1:2 PCA with MgCl_2_ .7H_2_O for 48 h at 37°C. Controls consisted of cells exposed to heat inactivated human serum (adapted from (62). Experiments were performed in quadruplicate.

### Complement binding

*P. aeruginosa* regular and persister cells were collected by centrifugation, resuspended in 0.85% saline to a concentration of 10^7^ cells/mL. The saline was supplemented with 100 μL of PBS or 100 μL of complete human serum to a final concentration of 10%. Cultures were incubated for 30 min to allow for binding of complement proteins to the bacterial cells, after which, the complement activity was stopped with 10 mM EDTA. Cells were then washed by centrifugation (16,000 *x g* for 4 min at 4°C), unbound proteins from the serum were decanted, and cells were resuspended in PBS. Serial dilutions of each population were performed, and each dilution was stained. Bacterial cells were stained with BacLight Red (6 ng - Thermo Fischer, Waltham, MA, USA), and complement protein C3b was labeled with fluorescent Anti-C3 antibody (35 μg MP Bio). Samples were imaged using an epifluorescence microscope (Olympus BX46) at 100x magnification. Experiments were performed in quadruplicate with 5 images being taken per sample. Images were analyzed using Intensity Luminance V1 software (63). To assess the binding of C5b, regular cells and persister cells of *P. aeruginosa* were collected by centrifugation, washed in 0.85% saline three times by centrifugation (16,000 *x g* for 5 min at 4°C) then resuspended in 0.85% saline at a final concentration of 10^7^ cells/mL, and subsequently incubated in either 90% complete human serum or PBS for a period of 1.5, 3, or 24 hours. The complement reaction was then stopped with 10mM EDTA, and the cells were washed in 0.85% saline three times by centrifugation (16,000 *x g* for 5 min at 4°C). The cell pellets were resuspended in PBS and then ELISA assays were executed per manufacturer’s instructions to measure membrane bound C5b (Novus Biologicals).

### Maintenance and differentiation of THP-1 macrophages

THP-1 monocytes were cultured in suspension on RPMI 1640 complemented with 10%Fetal Bovine Serum, with media changes every 2-3 days. Cultures were split once they reached 8 x 10^5^ cells/mL. THP-1 monocytes were seeded at a concentration of 5 x 10^5^ cells/mL onto 24-well plates and differentiated into M0 Macrophages via the introduction of 100 nM phorbol 12-myristate 13-acetate (PMA) for 3 days at 37°C 5% CO_2_, after which they were ready for the experimental procedures (27, 64).

### Infection of THP-1 Macrophages

*P. aeruginosa* persister and regular vegetative cells were isolated and resuspended in infection media (27, 64) and standardized to 5 x 10^6^ cells/mL. Infections were initiated with a MOI of 10:1 and incubated for different time periods, including, 30, 60, 90, and 180 min. Following the infection period, THP-1 macrophages were washed twice with PBS and subsequently exposed to gentamycin (40 mg/L) for 1 h to remove any remaining extracellular bacteria (27, 64). Macrophages were then lysed with 10% Triton X-100 for 45 min and intracellular bacteria viability was quantified as described above. Experiments were performed in quadruplicate.

### Opsonization and Engulfment of *P. aeruginosa* persister cells

*P. aeruginosa* regular vegetative cells and persister cells, were incubated in a solution of 10% human serum or PBS for a period of 30 min(62). The complement reaction was stopped with 10mM EDTA, cells were washed via centrifugation (16,000 *x g* for 5 min at 4°C), followed by resuspension in PBS, and subsequently used to infect THP-1 macrophages as described above. Experiments were performed in quadruplicate.

### Elimination Assays of Intracellular bacteria

To assess the elimination of intracellular bacteria, infections were performed for a period of 90 min as described above. Once infections were stopped, and following the gentamicin exposure, infection media was replenished, and the cultures were incubated for further 24 h. After incubation, macrophages were lysed with 10% Triton X-100 for 45 min and intracellular bacteria viability was quantified as described above. Experiments were performed in quadruplicate (65).

### Flow Cytometry

To assess the size of persister cells relative to regular vegetative cells, persister cells were isolated as above, stained with BacLight Red (6 ng - Thermo Fischer, Waltham, MA, USA) and then fixed with paraformaldehyde. Flow Cytometry was performed using the BD Accuri C6 Plus system and the data were analyzed with the flow cytometry software FlowJo. Similarly, to assess the elimination of persister cells, THP-1 macrophages were infected as above with pre-stained bacteria for 90 min or 24 h, followed by fixation with paraformaldehyde and flow cytometry analysis as above. To determine macrophage polarization, THP-1 macrophages were infected as above with either regular or persister cells for 90 min followed by immunostaining for M1/M2 cell-surface marker proteins CD80 (Phycoerythrin (PE) ANTI-HU CD80, from Biolegend), CD86 (Phycoerythrin (PE) ANTI-HU CD86, from Biolegend), and CD206 (Allophycocyanin (APC) ANTI-HU CD206, from Biolegend), and DAPI (Thermo Fisher) DNA staining for 30 min. The cells were then fixed with paraformaldehyde and flow cytometry analysis was performed with the Bio-Rad ZE5 Cell Analyzer and FlowJo. Experiments were performed in quadruplicate.

### Quantitative Reverse Transcriptase PCR (qRT-PCR)

Relative transcriptional levels of THP-1 innate immune genes and engulfed *P. aeruginosa* 16s rRNA were quantified. To quantify the relative expression, infections were performed as described above. At the end of the incubation with gentamicin, TRIzol reagent was added to macrophages, the contents of each well were collected, and RNA was isolated using the Zymo RNA purification kit (Zymo Research). A total of 0.5 μg of RNA was used for cDNA synthesis and cDNA was generated using QScript cDNA Synthesis kit. Quantitative reverse transcriptase PCR (qRT-PCR) was performed with an Eppendorf Mastercycler Ep Realplex instrument (Eppendorf AG, Hamburg, Germany) and the Kapa SYBR Fast qPCR kit (Kapa Biosystems, Woburn, MA) with the oligonucleotides for THP-1 cells (obtained from Qiagen) and *P. aeruginosa* 16s rRNA (12). No template controls (NTC) and no reverse transcriptase (NRT) reactions were executed to confirm the lack of DNA contaminants during sample and mastermix preparation. Relative transcript quantitation was accomplished using Ep Realplex software (Eppendorf AG), with the transcript abundance (based on the threshold cycle [CT] value) being normalized to the housekeepers *gadph* for THP-1 and *cysD* (FW: CTGGACATCTGGCAATACAT; RV: TCTCTTCGTCAGAGAGATGC) for *P. aeruginosa* before the determination of transcript abundance ratios. Single-product amplification verification was accomplished through analysis of the melting curves. Experiments were performed at least in quadruplicate (66).

### ELISA assays of secreted cytokine

Macrophages were infected with *P. aeruginosa* regular and persister cells and the supernatant was collected at 0.5, 1.5, and 24 h of infection. The supernatant was centrifuged for 5 min at 16,000 *x g* to remove bacterial cells, and the resulting solution was assessed for the presence of cytokines. Samples were diluted up to 1:100, and ELISA assays were performed to quantify protein concentration, per manufacturer’s instructions using the following kits (Invitrogen, Carlsbad, CA, USA): IL-10 (BMS2152), TNF-α (BMS223HS), IL-6 (BMS213HS), and CXCL-8 (KHC0081). At the end of each assay, the absorbance of each sample was determined at 450 nm (DTX880 multimode detector, Beckman Coulter, CA). Cytokine concentrations were determined using standard curves generated in each assay, then accounting for the dilution factor. Experiments were performed at least in quadruplicate.

### Statistical analysis

All data were analyzed using GraphPad Prism 9.3.1. One-way ANOVA was performed for multivariant analysis followed by Tukey’s or Dunnett’s multiple comparison tests.

